# In differentiated HL-60 neutrophil-like cells, MRP1- (ABCC1-) mediated glutathione efflux stimulated by BzATP and P2X_7_ receptor signalling regulates exosome release through nSMase activity

**DOI:** 10.64898/2026.02.26.708162

**Authors:** Erica Muriana Tintor, Anfal Sharif, Lúcia Moreno-Sάnchez, Samuel Antwi-Baffour, Jameel M. Inal

## Abstract

Exosomes are endosome-derived extracellular vesicles (EVs) playing key roles in immune regulation and inflammatory signalling, yet with poorly defined mechanisms of biogenesis. We investigated how purinergic signalling and redox regulation intersect, controlling exosome formation in neutrophil-like cells (NLCs) differentiated from the HL-60 cell line. Differentiated NLCs displayed hallmark neutrophil features, including increased CD11b expression, reduced CD71, enhanced phagocytosis, robust reactive oxygen species generation, and cell-cycle arrest. CD63-positive exosomes released from NLCs fulfilled MISEV 2023 criteria, exhibiting typical morphology, density, and size distribution.

Stimulation with the P2X_7_R agonist BzATP induced a rapid, dose-dependent increase in exosome release correlating strongly with depletion of intracellular glutathione (iGSH) and activation of neutral sphingomyelinase (nSMase). Pharmacological inhibition of nSMase abrogated BzATP-induced exosome release, implicating ceramide-dependent intraluminal vesicle formation. Mechanistically, BzATP activated PI3K/AKT signalling, leading to stimulation of the ABC transporter MRP1/ABCC1, enhanced efflux of GSH, and a concomitant rise in extracellular GSH. Inhibition of PI3K/AKT or MRP1 prevented iGSH efflux, reduced nSMase activity, and significantly attenuated exosome secretion.

These findings define a previously uncharacterized pathway in NLCs whereby extracellular ATP acts as a danger signal triggering PI3K/AKT–MRP1–dependent redox gating of nSMase activity, thereby driving rapid exosome biogenesis. This mechanism provides insight into how neutrophils translate inflammatory cues into vesicle-mediated communication and highlights potential therapeutic targets to modulate pathological neutrophil-driven inflammation.

## 1. Introduction

Extracellular vesicles (EVs) are membrane-bound nanostructures mediating intercellular communication by transferring bioactive molecules such as proteins, lipids, and nucleic acids between cells [1]. Among EV subtypes, exosomes originate from the endosomal system being formed through inward budding of endosomal membranes, generating intraluminal vesicles (ILVs) within multivesicular bodies (MVBs). Upon fusion of MVBs with the plasma membrane (PM), these ILVs are released as exosomes, in contrast with microvesicles (MVs), which bud directly from the PM [2].

Exosomes maintain cellular homeostasis and orchestrate physiological processes, while also contributing to pathological states such as infection, cancer, and autoimmune disorders [3]. Their ability to deliver diverse cargo—including lipids, proteins, and microRNAs—to recipient cells via receptor-mediated interactions or membrane fusion underscores their importance in immune regulation and disease progression. Beyond natural functions, exosomes have attracted interest as therapeutic vehicles for targeted delivery of drugs or genetic material [3]. To harness this potential, understanding molecular mechanisms governing exosome biogenesis, especially in neutrophils where this remains unclear, is essential for incorporating therapeutic payloads and modulating vesicle release.

Despite advances in EV biology and understanding of exosome biogenesis in macrophages [2], signalling pathways regulating ESCRT-independent exosome secretion in neutrophils remain incompletely defined [4]. Neutrophils, key effectors of innate immunity, express functional P2X_7_ receptors (P2X_7_R) on their PM which, upon activation by the agonist BzATP, induce calcium influx, potassium efflux, respiratory burst, and membrane pore formation [5]. Neutrophil-like cells differentiated from the HL-60 promyelocytic leukaemia line provide a robust experimental model to study these processes, closely mimicking primary neutrophil behaviour [6].

Previous studies in macrophages have implicated extracellular ATP (eATP) and P2X_7_R signalling in exosome biogenesis through activation of neutral sphingomyelinase (nSMase), an enzyme negatively regulated by intracellular glutathione (iGSH) [7,2]. Depletion of iGSH increases Reactive Oxygen Species (ROS). By causing lysosomal dysfunction that would otherwise limit MVB accumulation, MVB numbers and exosome secretion increase [8]. Increased extracellular GSH (eGSH) suggests active transport by ATP-binding cassette transporters, including MRP1/ABCC1. MRP1 likely contributes to this pathway, as its inhibition reduces both GSH efflux and exosome release.

Extracellular ATP is a danger signal for neutrophils, activating their P2 receptors, that bind ATP, ADP, and UTP [5]. The 1-3 mM concentrations of eATP found locally, released from damaged cells, are sufficient to activate P2X_7_Rs triggering cell death pathways, inflammation (IL-1β release) and ROS production in neutrophils In this study we used BzATP which has a higher affinity than ATP for P2X_7_Rs, at concentrations up to 300 μM, to ensure maximal P2X_7_R activation and downstream proinflammatory responses. This concentration also overwhelms the anti-inflammatory signalling triggered by P2Y_11_ receptors (which can be triggered by low μM concentrations of eATP). Neutrophil-like cells differentiated from the promyelocytic leukaemia cell line, HL-60, were used as they represent an excellent model, mimicking primary neutrophils.

Building on these insights, the current study aimed to elucidate how purinergic signalling and redox regulation intersect to control exosome biogenesis in NLCs. Specifically, the P2X_7_R-mediated activation of PI3K/Akt signalling was investigated, its influence on ABC transporter activation, and the downstream effects on nSMase activity and exosome formation. By advancing our understanding of neutrophil vesiculation, novel targets for therapeutic modulation of inflammatory responses can be identified.

## 2. Materials and Methods

### 2.1 Differentiation of HL-60 cells to Neutrophil-like cells (NLCs)

The human myeloid HL-60 cells were obtained from HPA and maintained in RPMI 1640 medium with 5% EV-depleted FBS (120,000 ×g/18h, then ultrafiltration (amicon, 100 kDa MWCO, 3,500 ×g/60 min) and 100 U/ml penicillin, 100 µg/ml streptomycin at 37 °C in a humidified atmosphere (5% CO_2_). The cells were split every 2-3 days. After maintaining the cells for 3 weeks, differentiation to NLCs was induced. HL-60, (10^5^ cells/ml) were treated with 1 µM all-trans retinoic acid (ATRA) and 1% dimethyl sulfoxide (DMSO) for 5 days. Once differentiated, cells no longer needed splitting and were used within a further 5 days.

### 2.2 Isolation of exosomes

Firstly small EVs were isolated by differential centrifugation. The NLC cell culture supernatant of BzATP-treated cells (for 2h) was centrifuged (160 ×g/10 min/4 °C). The supernatant was centrifuged (3,000 ×g /30 min/4 °C) and new supernatant filtered through a 0.22 μm pore size filter before centrifugation (25,000 ×g /30 min/4 °C) and final centrifugation (100,000 ×g /60 min/4 °C) with a final wash in 0.22 μm pore-filtered PBS, re-centrifugation and resuspension in a minimal volume. Exosomes were specifically isolated by immunoaffinity (IA) using the Tetraspanin Exo-Flow Capture kit (System Biosciences). This was carried out according to the manufacturer’s protocol. Essentially, the CD63-biotin antibody was coupled to the magnetic streptavidin beads (CD63-MBs). These CD63-MBs were then used to collect exosomes (or CD63-positive EVs) from the small EVs (150 μg) isolated by differential ultracentrifugation/ultrafiltration. After washing off unbound material, exosomes were eluted with exosome elution buffer. Exosomal buoyant density was determined by OptiPrep density gradient centrifugation. By combining IA with nanosight tracking analysis (NTA) we were able to monitor exosome biogenesis in real-time, obviating the need for a reporter system.

### 2.3 Nanosight Tracking Analysis (NTA)

To measure EV sample concentration and distribution profiles of EV diameter, the NanoSight NS300system (Malvern Instruments Ltd, UK) was used as described previously [9]. CD63-enriched exosomes were collected from 3 different samples of NLCs, diluted 500-fold in 1 ml of DPBS and injected into the instrument. Videos (5 × 1 min) were recorded as per the manufacturer’s instructions (between 17.5 and 19.5 °C). The EV concentration and size profile was derived using the NanoSight Software NTA 3.2 (Malvern).

### 2.4 SDS-Polyacrylamide Gel Electrophoresis and western Blotting

Protein extraction and protein assays were carried out as described previously [10]. Essentially, NLC exosomal proteins were obtained by lysing exosomes in cold RIPA buffer (Sigma) containing protein inhibitors (Sigma) for 60 min on ice followed by centrifugation (16,250 ×g /20 min/4 °C). Protein concentrations were measured by BCA protein Assay or by NanoOrange (both Thermo Scientific) as described previously [11]. For SDS-PAGE, samples were denatured in β-mercaptoethanol (95 °C/10 min); for deglycosylation, SDS was neutralised with NP-40 and 10 μg protein typically deglycosylated with 500 units of PNGaseF (37 °C/16 h). Western blotting was carried out using the Mini-PROTEAN tetra system on 4-20% TGX gels (Bio-Rad) as outlined previously [9] and transferred to nitrocellulose by semi-dry electrotransfer. The mouse antibodies used to probe the membrane were: anti-CD63 (1/1,000; Abcam, ab271286), anti-LAMP-2 (1/500; Abcam; ab25631), anti-Alix (1/1,000; Abcam; ab117600); anti-GM130 (1/500; Sigma-Aldrich, MABT1363). A rabbit polyclonal anti-P2X_7_R recognising an extracellular domain was from Alomone labs (1/500; APR-008) and anti-β-actin (1/1,000; Abcam, ab8226). The secondary antibodies used were anti-mouse and anti-rabbit HRP-conjugated antibodies (Bio-Rad) used at a 1/3,000 dilution, the blot then being visualised using chemiluminescence (ECL) and the UVP digital imaging system.

### 2.5 Immuno-Transmission Electron Microscopy

Paraformaldehyde-fixed exosomes were labelled with immunogold by first placing on a formvar carbon-coated nickel grid. They were blocked in 5% BSA/PBST and then incubated for 1h in with mouse monoclonal against human CD9 used at a 1/100 dilution (BD Pharmingen, Clone M-L13). Six 10 minute washes was followed by incubation in the secondary antibody (5 nm gold particle anti-mouse antibody (1/50 dilution). The grid was washed in PBS, distilled water and negative staining carried out with 2% uranyl acetate (Agar Scientific, UK). The grids were visualised using a JEM-1200 EX II electron microscope (JEOL, Peabody, MA), digital images, up to 40,000 magnification being obtained on an AMT digital camera.

### 2.6 Phagocytosis assay

A flow cytometry phagocytosis assay was carried out with NLCs, using the pHrodo Green E. coli BioParticles Phagocytosis Kit for Flow Cytometry (Invitrogen) according to the manufacturer’s instructions. Essentially pHrodo Green E. coli BioParticles Conjugate were incubated for 1 h at 37 °C. To prevent uptake, reactions were placed on ice. Cells were harvested and stained with 7-AAD, CD45-PE/Cy7 (Biolegend), and CD3-BV421 (Biolegend). Phagocytosis was determined as an increase in pHrodo Green fluorescence relative to that observed with control sample by flow cytometry using a BD FacsCanto II flow cytometer (BD Biosciences, USA), incubated on ice, and calculated as the percentage of pHrodo Green-positive cells. Relative to the negative control, increased pHrodo green fluorescence was taken as a positive phagocytosis.

### 2.7 Cell cycle analysis

HL-60 cell DNA was stained with propidium iodide (PI) as previously described using the Guava cell cycle assay [12]. Cells (10^5^ –10^6^) were centrifuged and mixed with 200 μl ice-cold 70% ethanol and left at -20 °C overnight. After washing in PBS they were resuspended in 200 μl Guava Cell Cycle Reagent (containing PI and other dyes) and after 30 min at room temperature in the dark loaded onto the Guava 8HT to differentiate cells according to DNA content (G0/G1, S and G2/M); analysis based on 10,000 events per sample, was with the ModFit software, version 5.0.

### 2.8 Intracellular GSH assay

To measure intracellular reduced glutathione (GSH), the Glutathione Competitive ELISA kit from Sigma (#CS0260) was used according to the manufacturer’s instructions. Essentially ∼1 × 10^6^ washed cells suspended in PBS were homogenised with 5-Sulfosalicylic Acid (SSA), further lysed by freeze-thaw and then centrifuged (10,000 ×g/10 min), supernatants being assayed immediately (in triplicate) in a microtiter plate. Using the provided standards, a standard curve was made to calculate the GSH concentration which was normalised by protein amount (extracted from an equal number of cells).

### 2.9 Extracellular GSH assay

Extracellular reduced glutathione was measured using the GSH-GLO− Glutathione assay (Promega). The growth medium of the NLCs (lacking cysteine for the incubation period) was treated as the sample. It was clarified (700 ×g/5 min/4 °C). To 50 μl in the wells of a 96-well plate (kept on ice), an equal volume of 2x GSH-Glo reagent containing GSH and the luciferin derivative was added with incubation for 30 min in the dark followed by 15 min with luciferin detection reagent (100 μl). To eliminate media interference, blank wells were included (cell-free medium alone) to subtract background. Luminescence was recorded. A standard curve (1-15 μM GSH) was used to convert relative light units to GSH concentration.

### 2.10 nSMase Enzyme Activity Assay

Neutral sphingomyelinase (nSMAse) activity in NLCs was measured using the Neutral Sphingomyelinase (nSMase) activity kit from Echelon Biosciences Inc. (cat no. K-1800); the coupled enzymatic reactions starting with nSMase hydrolysing sphingomyelin to phosphorylcholine and ceramide, eventually producing a blue product measured at 595 nm. The assay was carried out according to the manufacturer’s instructions. Essentially, 100 μl of sample or Sphingomyelinase standards were added to the 96-well plate, together with an equal volume of the Reaction mixture, incubated at 37 °C for 4h and then read on a FLUOStar Omega plate reader at 595 nm. The standard curve of A_595_ against Sphingomyelinase activity (mU/ml) was used to calculate the activity in the experimental samples. The rate of substrate activity in mU/ml, as a measure of nSMase activity, was converted to specific activity in mU/mg using the total protein determination, in mg/ml (BCA assay). The specific values for the samples treated with BzATP (for 2h) at various concentrations were averaged as were the control, untreated samples. The percentage of basal activity was calculated using:

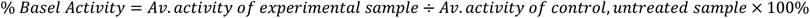

### 2.11 Measurement of intracellular ROS

HL-60 cells treated with 0.2 μM PMA for 60 min were assessed for intracellular ROS formation by staining with non-fluorescent 2’,7’-dichlorodihydrofluorescein diacetate (H_2_DCFDA, Santa Cruz). Samples with ∼5 × 10^5^ cells suspended in H_2_DCFDA (200 μM in PBS) were then incubated for 30 min in the dark at 37 °C and after washing in PBS, were analysed by flow cytometry. Inside the cells, H2DCFDA is deacylated by esterases and then oxidised by ROS converting it into the fluorescent DCF with an λ_max_ of 530 nm. This was measured with a BD FACSCanto II flow cytometer (BD Biosciences, USA) acquiring 10,000 events per sample. Differences between untreated and PMA-treated groups was tested using the non-parametric Mann-Whitney U-test for 2-sample comparisons of MFI (median fluorescence intensity).

### 2.12 Assay for intracellular calcium

This was carried out as described previously [11]. In brief, NLCs (5 × 10^4^ cells/well) were seeded into black-walled 96-well plates. After exposure to LPS (300 ng/ml for 3h) cells were washed and resuspended in phosphate buffered saline (PBS) loaded with 2 μM Fura 2-AM and 2 mM probenecid (30 min/37 °C. After washing off the loading solution and resuspending in PBS, steady state readings were taken on a fluorescent plate reader (λ_ex_340/380 nm, λ_em_540 nm) for 10 min. Upon addition of BzATP (300 μM) readings were measured for 50 min. Maximum fluorescence readings were obtained by lysing cells in 1% Triton X-100.

### 2.13 Flow cytometric analysis of HL-60 cells and human peripheral neutrophils

Flow cytometry was carried out as described previously [11]. Briefly, HL-60 cells, differentiated or not, were incubated in Fc receptor blocking solution for 20 min at room temperature before staining with anti-CD71 and anti-CD11b. Cells (5×10^5^ – 10^6^ cells/mL) washed twice in PBS, were incubated for 1h, in the dark, with FITC-labelled anti-CD71 (MEM-75, Thermo Fisher) and FITC-labelled mouse anti-human CD11b antibody (clone ICRF44; Bio-Rad). After washing analysis was with the Guava EasyCyte 8HT flow cytometer.

### 2.14 Statistical analyses

Data are presented as mean ± SEM, p<0.05 considered statistically significant. Statistical analyses used GraphPad Prism (version 10.6.1 for Windows). To evaluate the effects of BzATP on iGSH and EVs, a two-way analysis of variance (ANOVA) was conducted with Tukey’s post-hoc test. The relationship between BzATP and nSMAse activity was assessed using simple linear regression. For BzATP and nSMAse activity, dose-response data were fitted to a four-parameter logistic (4PL) curve using nonlinear regression with the least-squares method to determine IC50 values.

## 3. Results

### 3.1 Characterisation of neutrophil-like cells (NLCs) differentiated from promyelocytic HL-60 cells

To study exosome biogenesis in neutrophils, and to overcome problems with donor variability and short lifespan, rather than primary neutrophils, NLCs were chosen to ensure stable populations. To differentiate into NLCs, HL-60 cells were treated with ATRA plus DMSO for 3 and 5 days and showed increased expression of CD11b and decreasing expression of CD71 (Fig. 1A). The NLCs had increased capacity to phagocytose pHrodo Green E. coli particles (Fig. 1B and representative histograms in Fig. 1C). Upon PMA treatment, there was greatly increased ROS in NLCs compared to undifferentiated HL-60 (Fig. 1D and E) and decrease in S-phase (Fig. 1F). The NLCs were also confirmed to express P2X_7_R at increased levels compared to the undifferentiated HL-60 (Fig. 1G) and also increased expression when LPS-stimulated (Fig. 1H). To demonstrate their functionality, LPS-primed NLCs treated with BzATP resulted in an increase in Ca^2+^ influx, which could be inhibited with the P2X_7_R antagonist, A740003 (Fig. 1I).

**Fig. 1.**
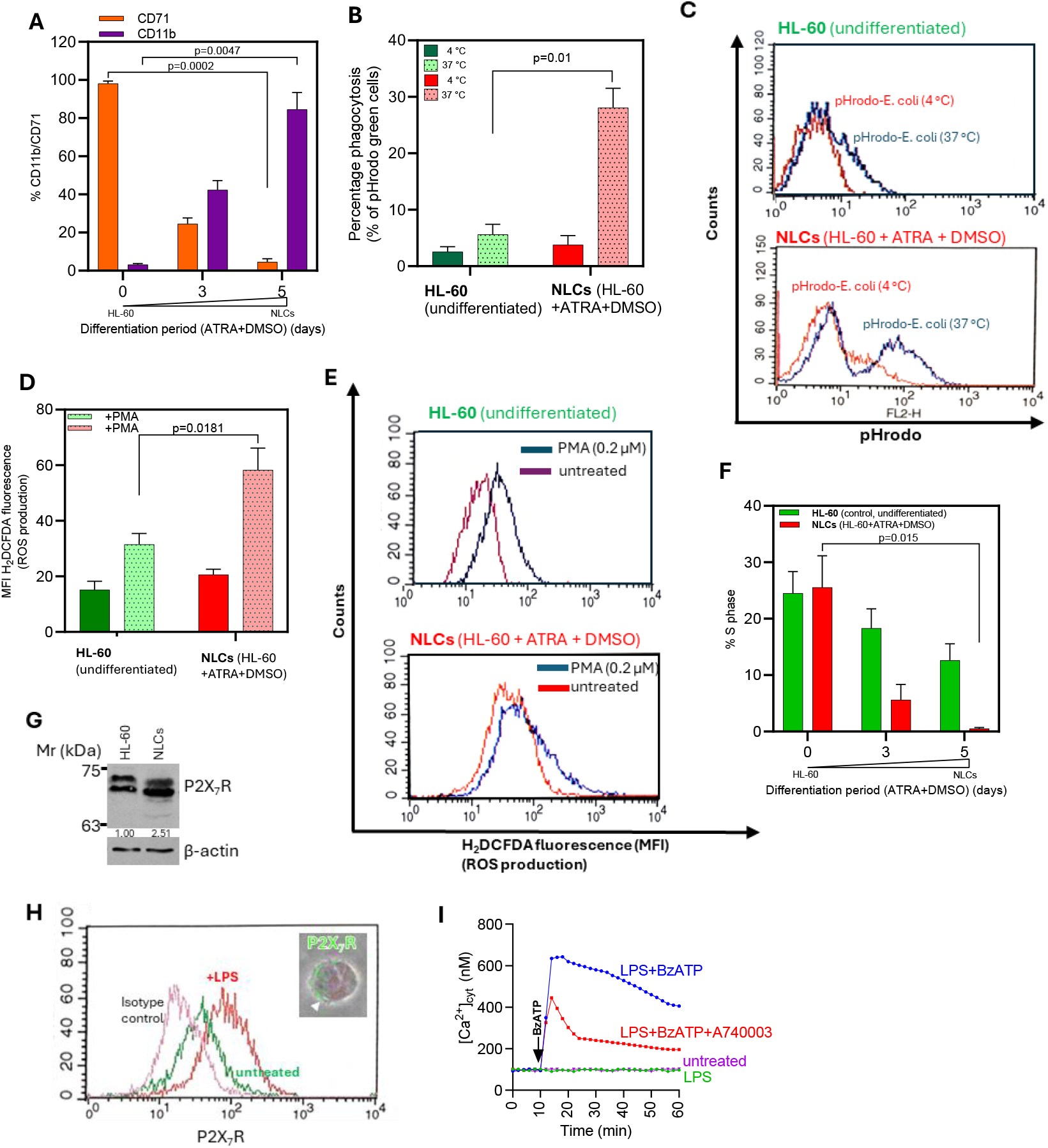
(A) Flow cytometry for expression of CD71 and CD11b was carried out after 0, 3 and 5 days of treatment of HL-60 cells with ATRA + DMSO using a Guava EasyCyte 8HT flow cytometer. (B), Percentage phagocytosis of pHrodo Green E. coli particles by NLCs (HL-60 cells differentiated for 5 days with ATRA + DMSO) compared to undifferentiated HL-60 cells, was measured using the FACSCanto II flow cytometer, representative histograms of engulfed particles measured as pHrodo green fluorescence intensity shown in (C). ROS production was measured as H2DCFDA fluorescence (MFI) on HL-60 and NLCs upon PMA stimulation (D) with representative histograms in (E). Cell cycle analysis (% S phase) for differentiated HL-60 after 3 and 5 days (F) and expression of P2X_7_R in HL-60 and NLCs by western blotting (G) showing band intensities normalised to β-actin, made relative to undifferentiated control, and by flow cytometry (H) and immunofluorescence microscopy (inset, H). Ca^2+^ uptake was monitored using Fura 2-AM upon adding BzATP to LPs-stimulated NLCs (I). Data are presented as mean ± SEM. Statistical significance was determined using an unpaired, two-tailed Student’s t-test (n=5 per group).

### 3.2 BzATP stimulation of NLCs and associated reduction of iGSH coupled with increased nSMAse activity and exosomal release

The CD63-enriched EVs (henceforth called exosomes) were isolated from NLCs by differential centrifugation and immunocapture using biotinylated anti-CD63 capture antibodies conjugated to streptavidin-conjugated magnetic beads. They were visualised by immuno-gold transmission electron microscopy using anti-CD9 (Fig. 2A). This revealed CD63-expressing “cup-shaped” EVs (a typical, though artefactual morphology of exosomes seen in TEM). NTA gave a peak modal size of 112 nm and range of 30-168 nm (Fig. 2B). Their buoyant density was 1.10-1.12 g/ml (Fig. 2C). For CD63, the NLCs (differentiated HL-60) showed hyperglycosylation (and increased expression of deglycosylated CD63) compared to undifferentiated HL-60 cells (Fig. 2D). The exosomes released from these cells were confirmed positive for EV markers (CD63 and Alix) but not the negative control Golgi apparatus marker, GM130 (Fig. 2E). Purity was confirmed with a total protein: particle number ratio <10^10^ particles/μg protein.

**Fig. 2.**
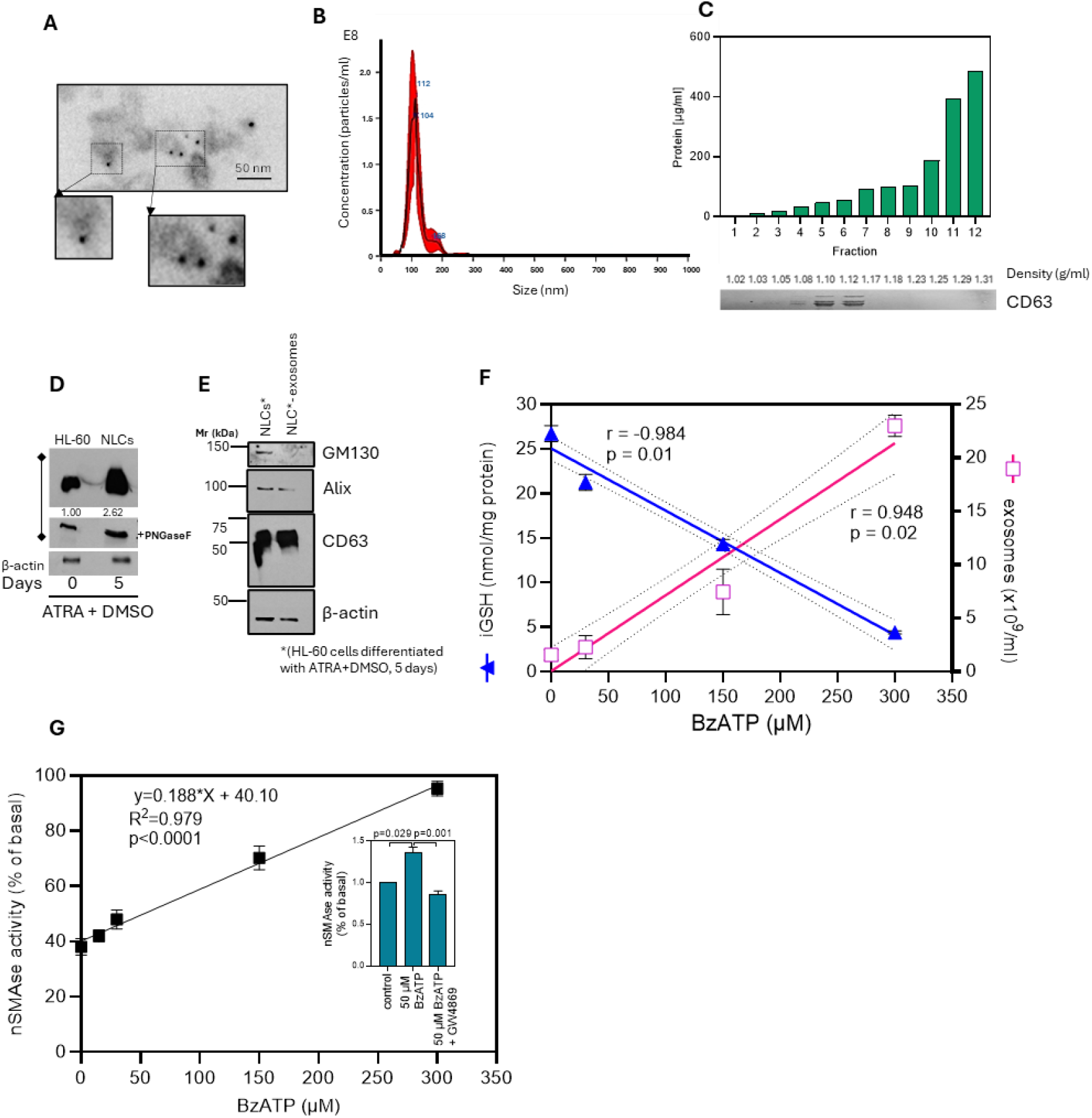
Immuno-transmission electron microscopy was used to reveal isolated CD63-positive exosomes, using anti-CD9 and 5 nm immunogold-conjugated secondary antibody (A) and their size range and peak modal size measured by NTA (B). Optiprep density gradient centrifugation showed the exosomes to have a density of 1.10-1.12 g/ml (C). Western analysis showed 5-day differentiated HL-60 cells (NLCs) to have increased expression of CD63 (D); image J 1.52p was used to calculate fold-changes of deglycosylated CD63 expression, band intensities normalised to β-actin and compared relative to control. The exosomal expression of EV markers was also compared with their parental cells (E). NLCs exposed to increasing BzATP concentrations up to 300 μM, were assayed for iGSH and concentration of released exosomes (F) and nSMAse activity (G) and in the presence of nSMAse inhibitor, GW4869 (F, inset). For (D) and (E) bars represent the mean ± SEM of n=3 independent biological replicates. Relationship between BzATP and iGSH, exosomes and nSMAse activity shown by linear regression fit, statistical significance of correlation indicated by R^2^ or r value and associated p-value.

Upon treatment of NLCs for 30 min with increasing concentrations of BzATP (eATP being a damage-associated molecular pattern (DAMP)), a strong positive correlation was found with exosome release (r=0.948, p=0.02) which was concomitant with a strong negative correlation (r=-0.984, p=0.01) with iGSH (Fig. 2F). Since iGSH is known to inhibit nSMAse [13] which is implicated in ILV formation in MVBs [14], we surmised that increasing BzATP (and therefore limiting iGSH) would lift inhibition of nSMAse, thus increasing its activity as demonstrated in Fig. 2G. To confirm nSMase-specific activity, we incorporated a specific nSMase inhibitor, GW4869, in NLCs similarly stimulated with BzATP (Fig. 2G (inset)).

### 3.3 iGSH is exported in a MK-571-inhibitable manner in response to BzATP resulting in a drop of iGSH and concomitant increase in exosomal release

BzATP-mediated P2X_7_R and P2Y_11_R signalling activates MRP1 through PI3K/AKT stimulation of aerobic glycolysis and production of ATP within minutes, which MRP1 needs, to actively pump substrates out of the cell [14]. We were able to demonstrate this in NLCs by the efflux of calcein-AM with increasing BzATP (Fig. 3A) and that this could be inhibited by MK-571 (Fig. 3A). The BzATP/PI3K/AKT-activated MRP1-mediated removal of iGSH from the cell should therefore result in increased eGSH. This was confirmed by demonstrating 150 μM BzATP significantly increasing eGSH, in a MK-571-inhibitable manner (Fig. 3C). We also found that two drugs affecting MRP1, MK-2206 which inhibits PI3K/AKT and therefore activation of MRP1 and MK-571 which blocks active MRP1, both inhibited EV release from BzATP-stimulated NLCs (Fig. 3C and E). To once more show decreasing iGSH with increasing concentrations of applied extracellular BzATP, it was then further showed that MK-571 could lessen this reduction in iGSH (Fig. 3D) and reduce the increase of exosome release (Fig. 3E).

**Fig. 3.**
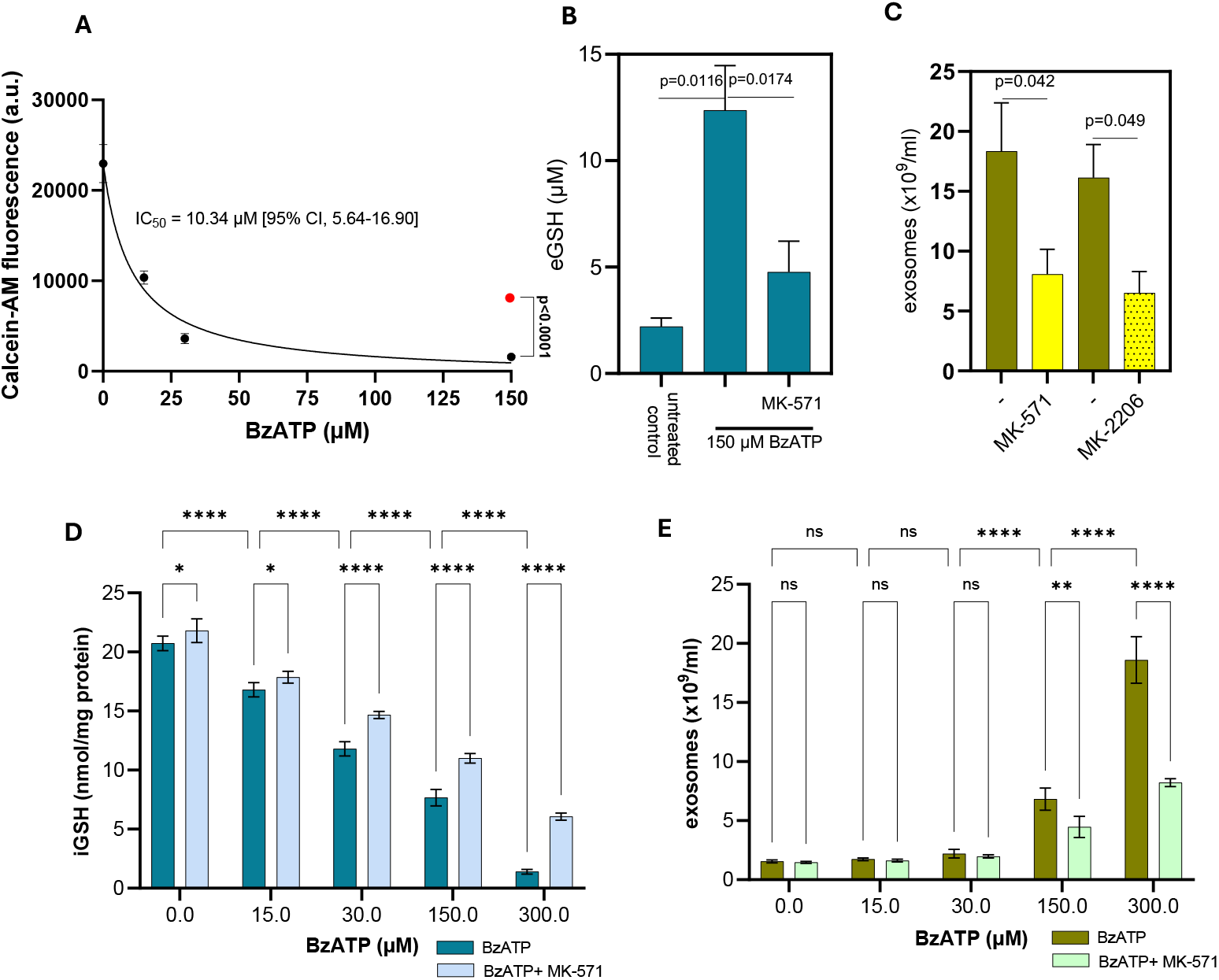
In (A), calcein-AM was measured in NLCs exposed to BzATP up to 150 μM, the filled red circle indicating the inhibitory effect of MK-571 on calcein-AM export. Increased eGSH (B) and exosome release (C) was measured upon BzATP stimulation of NLCs (also in the presence of inhibitors MK-571 and MK-2206). With increased expression of BzATP, iGSH (D) and exosome release (E) was assayed (also in the presence of MRP1-blocking MK-571) with increasing BzATP. Data are presented as mean ± SEM of n=3 independent biological replicates. Statistical significance was determined by 2-way ANOVA followed by Tukey’s post-hoc for pairwise comparisons between BzATP treated in the absence or presence of MK-571 and comparisons against the untreated control. Significance is indicated as *p < 0.05, **p < 0.01, and ***p < 0.001.

## 4. Discussion

Differentiation of HL-60 cells with ATRA+DMSO yielded NLCs exhibiting hallmark features of mature neutrophils including increased (CD11b) and decreased (CD71) abundance, enhanced phagocytic capacity, heightened PMA-induced ROS generation, and cell-cycle arrest. NLCs provide a robust surrogate for primary neutrophils in vesicle biogenesis studies. Exosomes purified from NLCs fulfilled ISEV-accepted criteria (MISEV 2023) [15] including CD63/Alix positivity with GM130 exclusion, “cup-shaped” morphology by immuno-TEM of CD63-positive exosomes, and a peak modal diameter near 112 nm by NTA.

### 4.1 Purinergic, redox, and sphingolipid coupling in exosome biogenesis

A key observation was the rapid, dose-dependent increase in exosome release after 30 min of BzATP treatment, tightly inversely correlated with iGSH levels. As reduced iGSH is a known endogenous inhibitor of nSMase, its depletion would be expected to disinhibit nSMase, increase ceramide production, and promote intraluminal vesicle (ILV) formation within MVBs, thereby driving exosome secretion [16,17,18,19]. Consistent with this model, pharmacological nSMase blockade with GW4869 curtailed BzATP-evoked vesicle release, implicating nSMase activity as a necessary step in the pathway [16].

Mechanistically, eATP (a prototypic DAMP) engages purinergic receptors, notably P2X_7_R (but also P2Y_11_ in neutrophils), to initiate signalling programs intersecting with lipid metabolic enzymes needed for vesicle biogenesis [20]. Our data place redox regulation at the centre of this cascade: BzATP stimulation lowers cytosolic iGSH while increasing eGSH, and the magnitude of exosome release positively tracks this GSH efflux. Because ceramide-dependent ILV formation is sensitive to the iGSH/nSMase axis, the ATP-driven exporter-mediated loss of iGSH functions as a rapid molecular switch to gate nSMase activity and exosome output.

### 4.2 Evidence for a PI3K/AKT–MRP1 (ABCC1) arm upstream of iGSH efflux

Upstream of nSMase, the transporter and signalling data converge on a PI3K/AKT–MRP1 (ABCC1) efflux mechanism. First, BzATP increased calcein-AM efflux, an observation consistent with MRP1 activity, and MK-571 (an MRP1 inhibitor) blocked this effect supporting the role of MRP1 in ATP-dependent efflux processes [21]. Second, MK-571 attenuated the BzATP-induced depletion of iGSH and the rise in eGSH [21]. Third, pharmacologic inhibition of AKT (MK-2206) or direct MRP1 blockade (MK-571) reduced exosome release. Our results showed in neutrophils, that BzATP activates P2X_7_R and PI3K/AKT signalling, in turn stimulating MRP1-mediated iGSH efflux and providing relief of nSMase inhibition. The subsequent accumulation of ceramide resulted in exosome biogenesis. While PI3K/AKT modulation of ABC transporters is well documented (particularly for ABCG2 localization and activity) and MRP1 is recognized for transporting iGSH and iGSH-conjugates, further genetic validation would strengthen the specific link to neutrophil-lineage cells.

### 4.3 Biological significance and potential translation

The delineated pathway underscores how NLCs can transduce an ATP danger signal into fast, post-translational control of vesicle biogenesis via redox gating. By exporting iGSH through MRP1, cells swiftly lift nSMase inhibition and trigger ceramide-dependent ILV budding, enabling rapid exosome release without requiring de novo gene expression. In inflammatory microenvironments enriched in eATP, such a mechanism could amplify intercellular communication through exosomal cargoes (proteins, lipids, miRNAs) and potentially modulate innate responses and tissue outcomes [22].

Neutrophil exosomes are a key element in hyperinflammatory disease. They transport inflammatory cargo (miR155, leukotriene B4) which further activates neutrophils and other immune cells and damaging enzymes (cathepsin G, MMP-9, proteinase 3 and neutrophil elastase) leading to organ dysfunction. This perpetuates chronic inflammation preventing tissue repair. Dangerous hyperinflammation with excessive neutrophil activity and NETosis (release of Neutrophil Extracellular Traps), is part of the pathology of various diseases including sepsis (severe infection), autoimmune diseases, chronic lung diseases (COPD) and certain cancers. From a translational perspective, pharmacological targeting of PI3K/AKT, MRP1, or nSMase offers testable strategies to dampen excessive neutrophil vesiculation in hyperinflammatory states, while redox-modulating agents (e.g., GSH donors) might suppress ATP-evoked exosome release [23].

### 4.4 Conclusion

Our findings summarised in Figure 4, support a framework in which BzATP-triggered purinergic signalling activates a PI3K/AKT–MRP1 axis to export iGSH, relieving nSMase inhibition and promoting ceramide-driven ILV formation and exosome release from NLCs. Going forward, the pathway’s pharmacological inhibition, and integration of damage-associated ATP with redox-sensitive lipid biogenesis highlight points for modulating neutrophil vesiculation in inflammation.

**Fig. 4.**
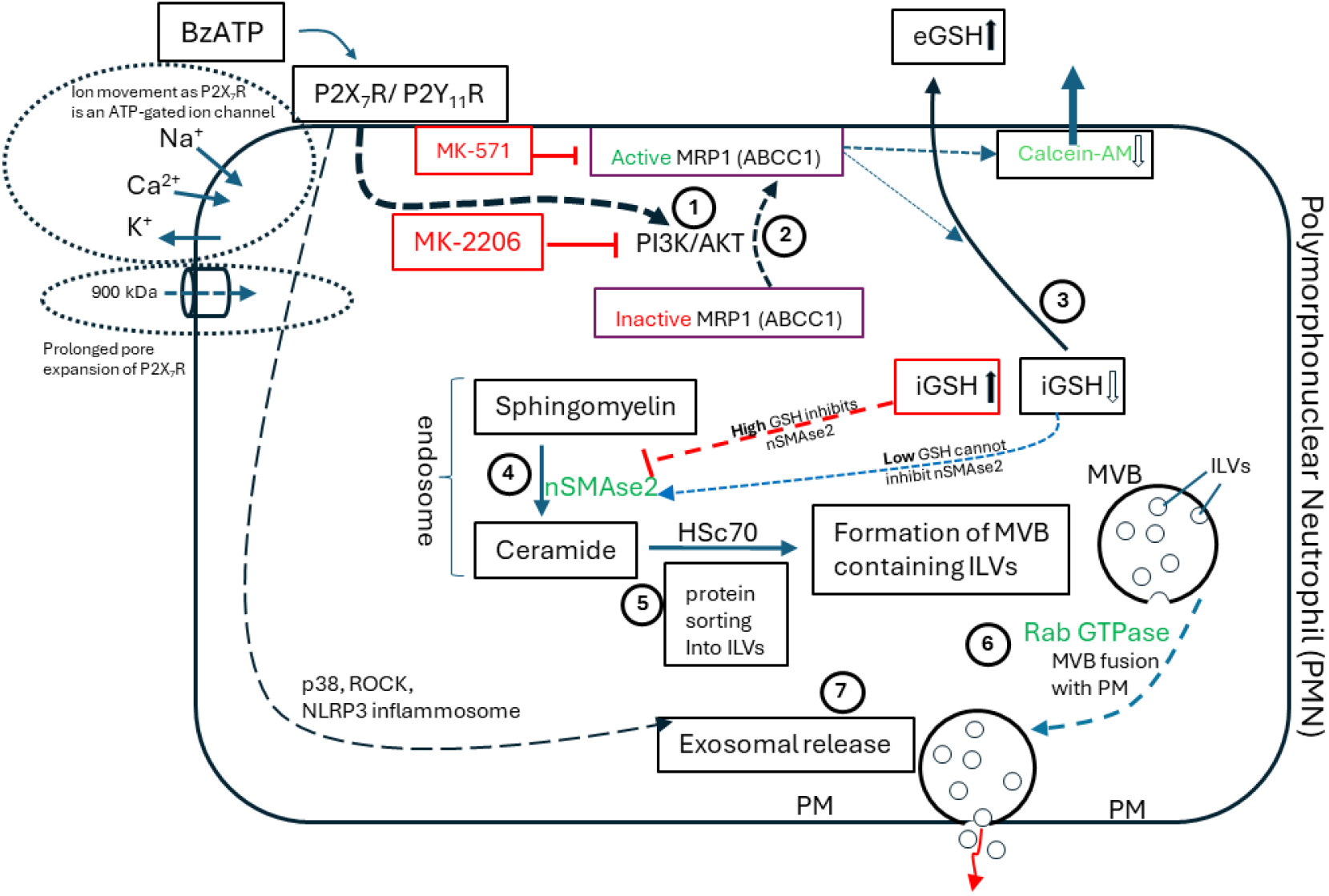
Model for MRP1- (ABCC1-) mediated glutathione efflux in neutrophil-like cells in which P2X_7_Rs stimulated by BzATP regulates nSMAse activity and exosome release.

## Acknowledgements

Royal Society Grant International Exchange IV0871706

